# A mechanistic model for reward prediction and extinction learning in the fruit fly

**DOI:** 10.1101/2020.12.03.409490

**Authors:** Magdalena Springer, Martin Paul Nawrot

**Author notes:** www.computational-systems-neuroscience.de.

## Abstract

Extinction learning, the ability to update previously learned information by integrating novel contradictory information, is a key mechanism for adapting our behavior and of high clinical relevance for therapeutic approaches to the modulation of maladaptive memories. Insect models have been instrumental in uncovering fundamental processes of memory formation and memory update. Recent experimental results in *Drosophila melanogaster* suggest that, after the behavioral extinction of a memory, two parallel but opposing memory traces coexist, residing at different sites within the mushroom body. Here we propose a minimalistic circuit model of the *Drosophila* mushroom body that supports classical appetitive and aversive conditioning and memory extinction. The model is tailored to the existing anatomical data and involves two circuit motives of central functional importance. It employs plastic synaptic connections between Kenyon cells and mushroom body output neurons (MBONs) in separate and mutually inhibiting appetitive and aversive learning pathways. Recurrent modulation of plasticity through projections from MBONs to reinforcement-mediating dopaminergic neurons implements a simple reward prediction mechanism. A distinct set of four MBONs encodes odor valence and predicts behavioral model output. Subjecting our model to learning and extinction protocols reproduced experimental results from recent behavioral and imaging studies. Simulating the experimental blocking of synaptic output of individual neurons or neuron groups in the model circuit confirmed experimental results and allowed formulation of testable predictions. In the temporal domain, our model achieves rapid learning with a step-like increase in the encoded odor value after a single pairing of the conditioned stimulus with a reward or punishment, facilitating single-trial learning.

## Introduction

Fruit flies can learn to associate an odor stimulus with a positive or negative consequence, e.g. food reward or electric shock punishment. In the training phase flies are typically exposed to two odors (differential conditioning) where one odor (conditioned stimulus minus, CS-) is perceived alone whereas a second odor (conditioned stimulus plus, CS+) is presented together with either reward or punishment (unconditioned stimulus, US). Once an association has formed between the CS+ and the respective US, the learned anticipation of the US can be observed in a memory test that enforces a binary choice behavior between the CS+ and the CS-(Tempel, Bonini, Dawson, & Quinn, 1983; Tully & Quinn, 1985). A single learning trial can be sufficient to form a stable memory in the fruit fly (Beck, Schroeder, & Davis, 2000; Krashes & Waddell, 2008; Zhao et al., 2019) and other insect species (see Discussion).

The prediction error theory (Rescorla & Wagner, 1972) describes a basic theoretical concept of classical conditioning. It assumes that the efficacy of learning is determined by the momentary discrepancy (or error) between the expected and the received reinforcement (i.e. reward or punishment). In vertebrates, it has been shown that prediction error coding dopaminergic neurons (DANs) are involved in learning (Schultz, 2016; Schultz, Dayan, & Montague, 1997). Recent studies give rise to the assumption that dopaminergic neurons could play a similar role in *Drosophila melanogaster* (Eichler et al., 2017; Eschbach et al., 2020; Felsenberg, Barnstedt, Cognigni, Lin, & Waddell, 2017; Felsenberg et al., 2018; Hammer, 1997; Ichinose et al., 2015; Riemensperger, Völler, Stock, Buchner, & Fiala, 2005) and other insects (Terao & Mizunami, 2017), rejuvenating an earlier hypothesis based on experimental observations in the honeybee (Hammer, 1997).

Re-exposing the flies to the CS+ after successful training and in the absence of the US leads to a reduction of the previously learned behavior (Felsenberg et al., 2017, 2018; Schwaerzel, Heisenberg, & Zars, 2002; Tempel et al., 1983). This new learning is called extinction learning and has been observed across invertebrate (Eisenhardt, 2014; Eisenhardt & Menzel, 2007) and vertebrate species (Bouton, 2004, 2017; Dudai, 2004; Myers & Davis, 2002; Pavlov, 1927; Tully & Quinn, 1985). Following prediction error theory, extinction learning is caused by the mismatch between the expected outcome (predicted US) based on the initial learning and the actual outcome (no US). In humans, extinction learning is of high clinical relevance and is applied in the treatment of humans suffering from maladaptive memories (Bouton, 2017; Chiamulera, Hinnenthal, Auber, & Cibin, 2014; Delamater & Westbrook, 2014; Quirk & Mueller, 2008; Walsh, Das, Saladin, & Kamboj, 2018). The majority of behavioral experiments in invertebrates and vertebrates suggest that extinction protocols, including the applied exposure therapy in humans, does not erase a memory but rather leads to the formation of a parallel but opposing memory, the extinction memory (Bouton, 2004; Eisenhardt & Menzel, 2007). Despite the relevance for clinical therapy, our understanding of the neural circuit mechanisms underlying extinction learning is still in its infancy.

We study a computational neural circuit model of the fruit fly mushroom body that captures the most recent anatomical and physiological facts. The model proposes detailed mechanisms for associative learning, extinction learning, prediction error coding, and single trial learning. We compare the model outcomes quantitatively to the results of recent behavioral and physiological studies and we derive novel experimental predictions by mimicking neurogenetic manipulations of relevant neuron groups.

## Results

### Circuit model of the *Drosophila* Mushroom Body

We implemented a neural network model of the olfactory memory circuit of *Drosophila melanogaster* (Figure 1) where individual neurons exhibit trial-resolved activation dynamics (see Methods). Our network model integrates experimentally confirmed connections in the adult fruit fly. It involves three feed-forward layers of the antennal lobe projection neurons (PN), the mushroom body (MB) Kenyon cells (KCs), and four individual mushroom body output neurons (MBONs) with specific lateral inhibitory connections. Reinforcing stimuli (US) of rewarding or punishing nature are mediated by two dopaminergic neurons (DANs). Each DAN receives specific feedback input from a single excitatory MBON and exerts a recurrent neuromodulatory effect on the plastic KC::MBON synapses.

**Figure 1:**
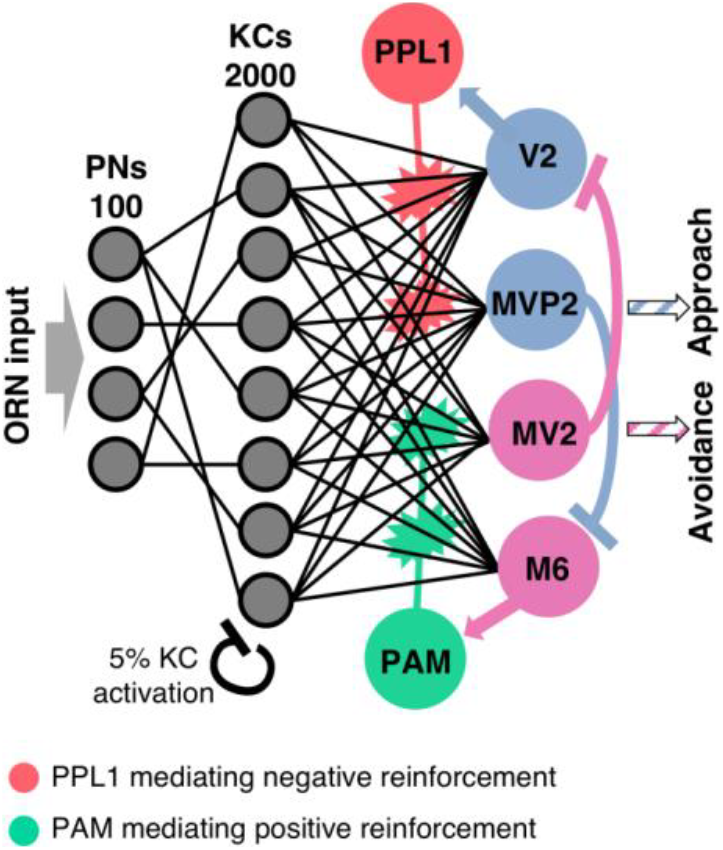
Circuit model. The olfactory pathway comprises three feedforward layers. Olfactory input activates an odor-specific combination of PNs. The connectivity matrix from PNs to the large population of KCs is divergent-convergent and sparse. The inhibitory feedback mechanism ensures population sparseness of 5% active cells in the KC layer. KCs are fully connected with four MBONs in the output layer. A positive or negative reinforcement stimulus directly excites the dopaminergic PAM or PPL1 neuron, respectively. Lateral inhibition between the MBONs and excitatory feedback from MBONs to DANs is crucial for the reward prediction and extinction mechanisms. Activation of the MVP2 output neuron mediates approach behavior, activation of MV2 mediates avoidance behavior. Behavioral preference towards an odor is calculated as the imbalance between the activations of MVP2 and MV2.

Stimulation with a particular odor is modeled as one specific input pattern activating 50 out of total 100 PNs, each with a random activation rate (see Methods). Similarity between different odors was defined as the percentage of overlap between the PN activation patterns (Figure 2C). This establishes a dense combinatorial odor code in the PN layer as reported experimentally for fruit flies (Olsen, Bhandawat, & Wilson, 2010; Wilson, Turner, & Laurent, 2004) and other species, notably the honeybee (Joerges, Küttner, Galizia, & Menzel, 1997; Krofczik, Menzel, & Nawrot, 2009), the locust (Broome, Jayaraman, & Laurent, 2006; Mazor & Laurent, 2005), and the moth (Namiki & Kanzaki, 2008).

**Figure 2:**
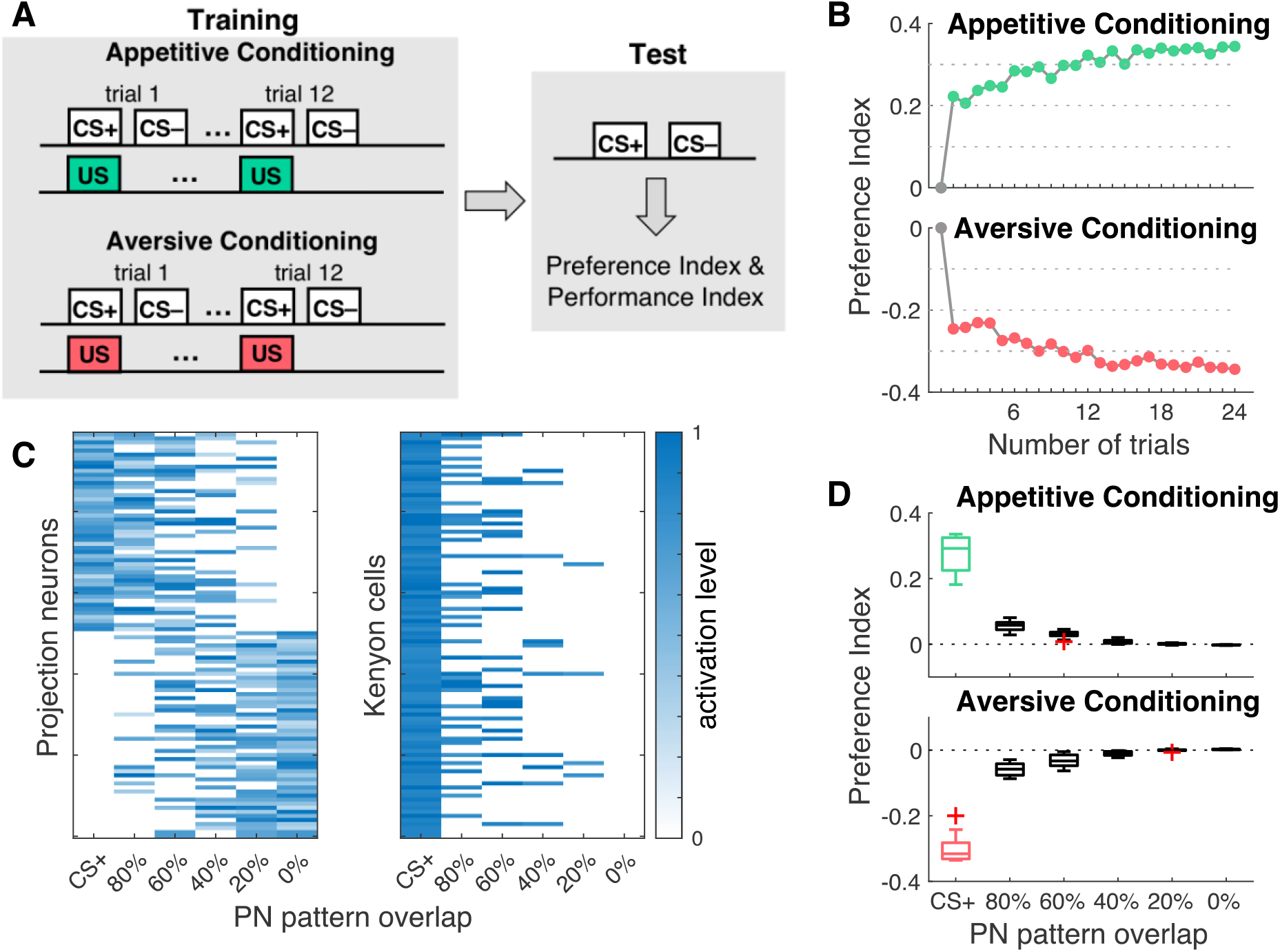
Differential conditioning leads to CS+ specific odor preference. **A)** Experimental protocols for simulation of associative conditioning experiments. The model was trained in a classical conditioning paradigm, in which a conditioned odor stimulus (CS+) was paired with a reward (green) or punishment (red). Subsequently, a second unconditioned odor stimulus (CS–) was presented without reinforcement. The standard protocol comprises 12 training trials. During the retention test, both odor stimuli were presented alone without reinforcement. Preference index and performance index in the model were computed after each trial and during the retention test. **B)** Dynamics of the odor preference index across 24 appetitive (top) and aversive (bottom) learning trials, respectively averaged across 10 networks. **C)** Left: Combinatorial response patterns in the PN population. Each odor stimulus activates 50% of all PNs. Similarities between the CS+ reference odor and five novel odors are defined by their overlap in the PN activation pattern (x-axis). Right: Activation patterns across the subpopulation of 100 KCs (5%) activated by the CS+ odor. Pattern overlap in the KC population reduces rapidly with decreasing odor similarity as expressed in the percentage of PN pattern overlap (x-axis). **D)** Generalization to different odors after associative conditioning to the CS+ odor. The model shows a significant CS+ approach (green) or avoidance (red) after 12 training trials as expressed in the preference index (n=15), which diminishes rapidly with decreasing odor similarity. Boxplots show the median and the lower and upper quartiles, whiskers indicate 1.5 times inter-quartile range, outliers are marked with ‘+’ symbol.

The connectivity between the PNs and KCs is divergent-convergent and random (Caron, Ruta, Abbott, & Axel, 2013) where each of the 2.000 KCs (Aso et al., 2009) connects to average 10 PNs (uniformly distributed, range of 5 - 15), matching the experimentally estimated numbers for *Drosophila* (Leiss, Groh, Butcher, Meinertzhagen, & Tavosanis, 2009; Turner, Bazhenov, & Laurent, 2008) and establishing a sparse connectivity for each of the postsynaptic KCs. In a second step, a threshold criterion was implemented such that only the 5% of KCs with the highest activation rates retain their activation (Figure 2C) while the activation rates of all other KCs was set to zero (Peng & Chittka, 2017). This enforces population sparse coding in the KC layer (Kloppenburg & Nawrot, 2014) as reported in physiological experiments in the fruit fly where, on average, ~5% of all KCs were activated by a single odor stimulus (Honegger, Campbell, & Turner, 2011; Lin, Bygrave, de Calignon, Lee, & Miesenböck, 2014; Turner et al., 2008).

In the MB output layer, we implemented four out of the 34 anatomically identified MBONs (Aso, Hattori, et al., 2014; Ito et al., 1998; Séjourne et al., 2011; Tanaka, Tanimoto, & Ito, 2008). These are MV2 (MBON-β1>α), M6 (MBON-γ5β′2a), MVP2 (MBON-γ1pedc>α/β) and V2 (MBON-α'1, MBON-α′3ap, MBON-α′3m, MBON-α2sc and MBON-α2p3p). The neuron cluster V2, consisting of five neurons, was modeled as a single neuron for simplicity. M6, MV2, MVP2 and V2 have been previously shown to be involved in odor valence coding with M6 and MV2 mediating odor driven avoidance behavior, and MVP2 and V2 promoting approach behavior towards an olfactory stimulus (Aso, Sitaraman, et al., 2014; Bouzaiane, Trannoy, Scheunemann, Plaçais, & Preat, 2015; Owald et al., 2015; Perisse et al., 2016; Séjourne et al., 2011; Ueoka, Hiroi, Abe, & Tabata, 2017). All KCs connect to each of the four MBONs (full connectivity, Figure 1). Our model includes two inhibitory lateral connections among MBONs.

The inhibitory synapses between MVP2 and M6 (Felsenberg et al., 2018; Perisse et al., 2016) have been suggested to be functionally relevant for aversive memory extinction (Felsenberg et al., 2018). We additionally assume a symmetric lateral inhibitory connection from MV2 to the V2 neuron. It has been shown that the MV2 neuron projects onto MBON-α2sc and MBON-α2p3p, which are part of the V2 cluster and it was hypothesized that the glutamatergic MV2 acts inhibitory on both neurons (Aso, Hattori, et al., 2014; Aso, Sitaraman, et al., 2014), as has been shown for glutamatergic neurons in the AL (W. W. Liu & Wilson, 2013).

### Reinforcing pathways and synaptic plasticity

In our circuit model the presence of the appetitive or the aversive US are signaled by two distinct neuromodulatory DANs, the PPL1 and PAM neuron, respectively (Figure 1). The single PPL1 neuron is representative of the PPL1 neuron cluster activated by aversive sensory stimuli such as electric shock (Aso et al., 2012, 2010; Claridge-Chang et al., 2009; Mao & Davis, 2009), the PAM neuron represents the PAM cluster that is activated by appetitive sensory stimuli such as sucrose (Burke et al., 2012; C. Liu et al., 2012; Waddell, 2013; Yamagata et al., 2015). Both neurons receive additional excitatory input from either the M6 or the V2 MBON (Eichler et al., 2017; Eschbach et al., 2020; Felsenberg et al., 2017, 2018; Ichinose et al., 2015).

The presence of a reinforcing signal (US) has differential effects on the DAN activation. A rewarding US leads to an excitation of PAM and can reduce excitation of PPL1 input (Eqn. 5a). Vice versa, a punishing reinforcer excites PPL1 and at the same time can modulate excitation of PAM input (Eqn. 5b). This modulatory effect in our model is based on an experimental study revealing that a negative reinforcement inhibits PAM-γ4 and PAM-γ5 neurons, whereas sugar feeding inhibits PPL1-γ2 and PAM-γ3 (Cohn, Ianessa, & Ruta, 2015).

Plasticity in our model exclusively resides in the KC::MBON synapses, reflecting the prevalent data-based hypothesis (see Discussion). These synapses are subject to plasticity according to the learning rule in Eqns. 7a and 7b. In the initial state of the model, the weights of all KC::MBON synapses are set to the same fixed value. In a given experimental trial, and for any synapse KC::MV2 or KC::M6, plasticity occurs when activation of the presynaptic KC coincides with activation of the reward-mediating PAM neuron. Likewise, any KC::V2 or KC::MVP2 synapse undergoes changes when the presynaptic KC and the punishment-mediating PPL1 neuron are active at the same time. The reduction of a synaptic weight is proportional to the DAN activation rate (for details see Methods). This establishes two distinct parallel but interconnected neuromodulatory pathways, each involving feedback from MBONs to the DANs that, in turn, can modulate KC::MBON synapses (Figure 1).

### Appetitive and aversive conditioning establish a behavioral odor preference

In a first set of experiments we subjected our model to a classical conditioning protocol. This protocol (Figure 2A) mimics standard training procedures in the fruit fly (see Methods). In each trial the trained odor, CS+, was paired with either reward or punishment (US). Subsequently a second odor stimulus was presented without reinforcement (CS-).

After a pre-defined number of training trials, we performed a memory retention test by presenting the model with the CS+ alone (no reinforcement) and subsequently with the unpaired odor (CS). We computed a preference index (Eqn. 8) based on the activation rates of approach and avoidance mediating MBONs for both odors, the CS+ and the CS-(see Methods). The model performance index (Eqn. 9) mimicks comparison with the behavioral performance index computed from experiments in the fruit fly.

Olfactory memories are established as a skew in the MBON network: aversive learning reduces CS+ driven responses in approach MBONs (MVP2-MBON and V2 cluster MBONs) resulting in a skew towards avoidance. In contrast, appetitive learning skews the network towards approach by reducing CS+ mediated input to avoidance coding M4 and M6 MBONs (Aso, Sitaraman, et al., 2014; Owald et al., 2015; Owald & Waddell, 2015; Perisse et al., 2016; Zhang, Noyes, Zeng, Li, & Davis, 2019).

We explored the preference index during the retention test for a varying number of training trials (Figure 2B). In the naïve state, i.e. before the first training trial, the model did not reveal a preference. This is due to the balanced synaptic weight initialization of the model. A single appetitive training trial yielded a strong preference towards the rewarded odor, the CS+. With each additional training trial, the preference index further incremented by a small amount and thus additional training led to only a gradual increase of the preference index saturating after ~10-15 trials. Aversive conditioning revealed a negative preference index reaching a similar absolute level after a single training trial (Figure 2B, lower panel). Both, single trial learning and saturation of behavioral performance within few trials matches the experimental observations during associative learning in insects (see Discussion).

For all subsequent experiments we chose 12 training trials, both for the training and the reactivation phase. This number mimics the standard experimental protocol for aversive conditioning in fruit flies, in which a group of animals is exposed to 12 subsequent electric shocks while constantly being exposed to the CS+ (Felsenberg et al., 2018; Scheunemann et al., 2013; Tully & Quinn, 1985).

### Generalization to novel odors

We next analyzed how the learned preference to the CS+ odor generalizes to novel odors of varying similarity with the CS+ odor by comparing the preference indices towards CS+ and towards a novel odor during the retention test (Figure 2D). We designed the pattern of activated PNs for each novel odor such that it has a defined overlap with the CS+ pattern, ranging from zero to 80% overlap (see Methods). Our results in Figure 2D predict that generalization to a novel odor is rather low even for the highest odor similarity and decays with decreasing odor similarity, effectively reaching zero generalization for odors that share 40% or less activated PNs with the CS+ odor. For the subsequent extinction learning experiments, we used CS-odors that had 60% pattern overlap with the CS+ odor.

### Extinction learning significantly reduces the conditioned odor response

We analyzed extinction learning after appetitive and aversive conditioning according to the stimulation protocol in Figure 3A. The initial training phase was followed by a reactivation phase during which the model is repeatedly presented with the CS+ odor alone (no reinforcement). This matches the experimental protocols for appetitive (Felsenberg et al., 2017) and aversive (Felsenberg et al., 2018) conditioning and subsequent extinction learning and allows for a direct comparison of our simulation results with the experimental results, both at the behavioral and the physiological level (cf. Table 1).

**Figure 3:**
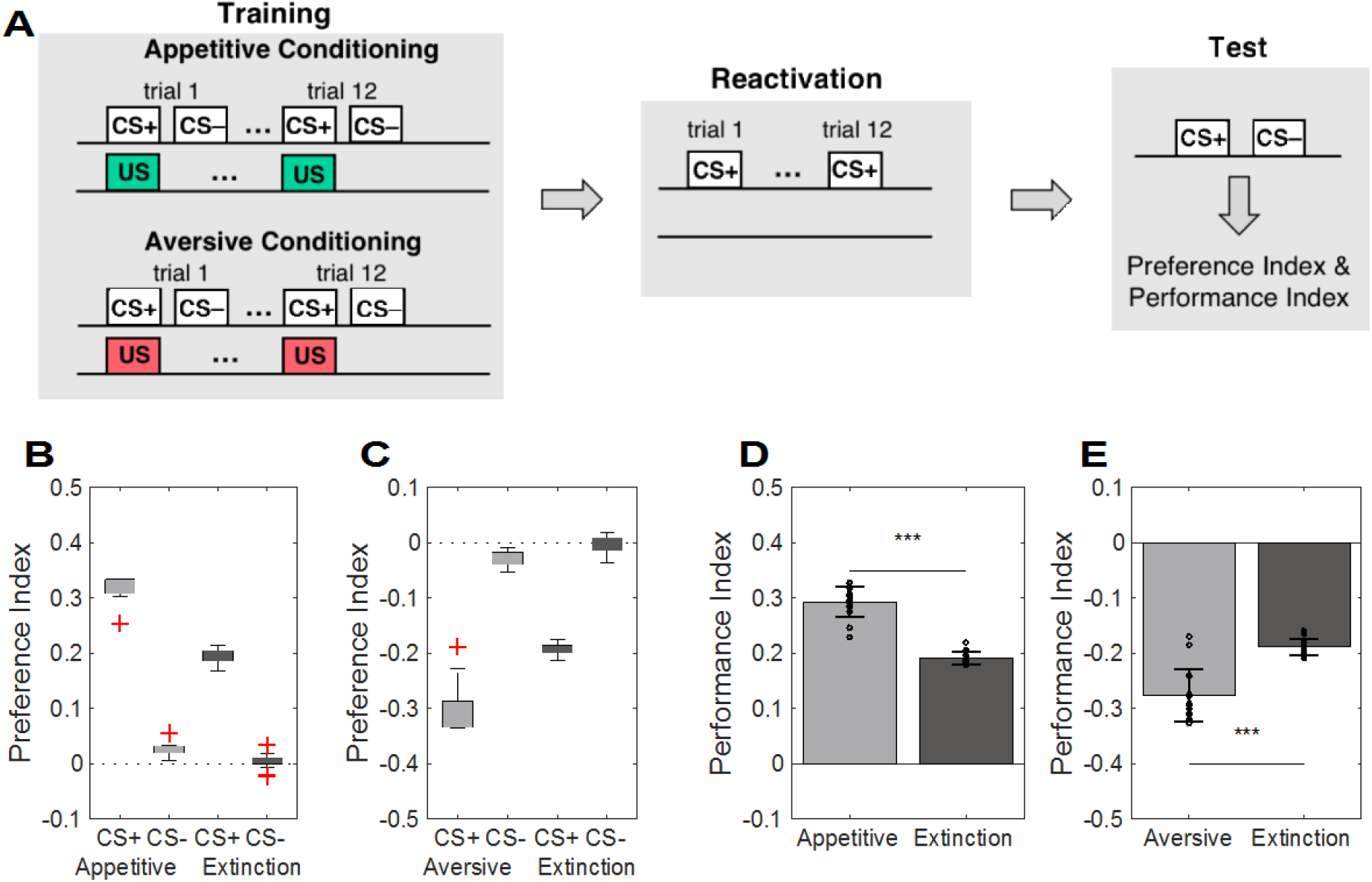
CS+ odor reactivation leads to memory extinction. **A)** Training and test protocol as in Fig 2A. In the extinction protocol, the training phase is followed by an odor reactivation phase, in which the CS+ was presented alone for 12 consecutive trials before performing the retention test. **B)** The appetitive conditioning protocol (light gray) induced a strong and significant preference for the CS+ odor and a weak and significant preference for the CS-odor. Extinction learning partly abolished the CS+ preference during the retention test (dark gray). The preference index for the CS+ odor was significantly lower than after initial appetitive conditioning, for the CS-odor no more significant preference was observed after extinction learning. **C)** Symmetric results were observed for aversive conditioning. Again, extinction learning significantly reduced the learned preference for the CS+ odor. **D, E)** The performance indices were significantly reduced after extinction compared to the initial appetitive and aversive memories. Data is presented mean ± STD; n=15 independent models.

**Table 1.**
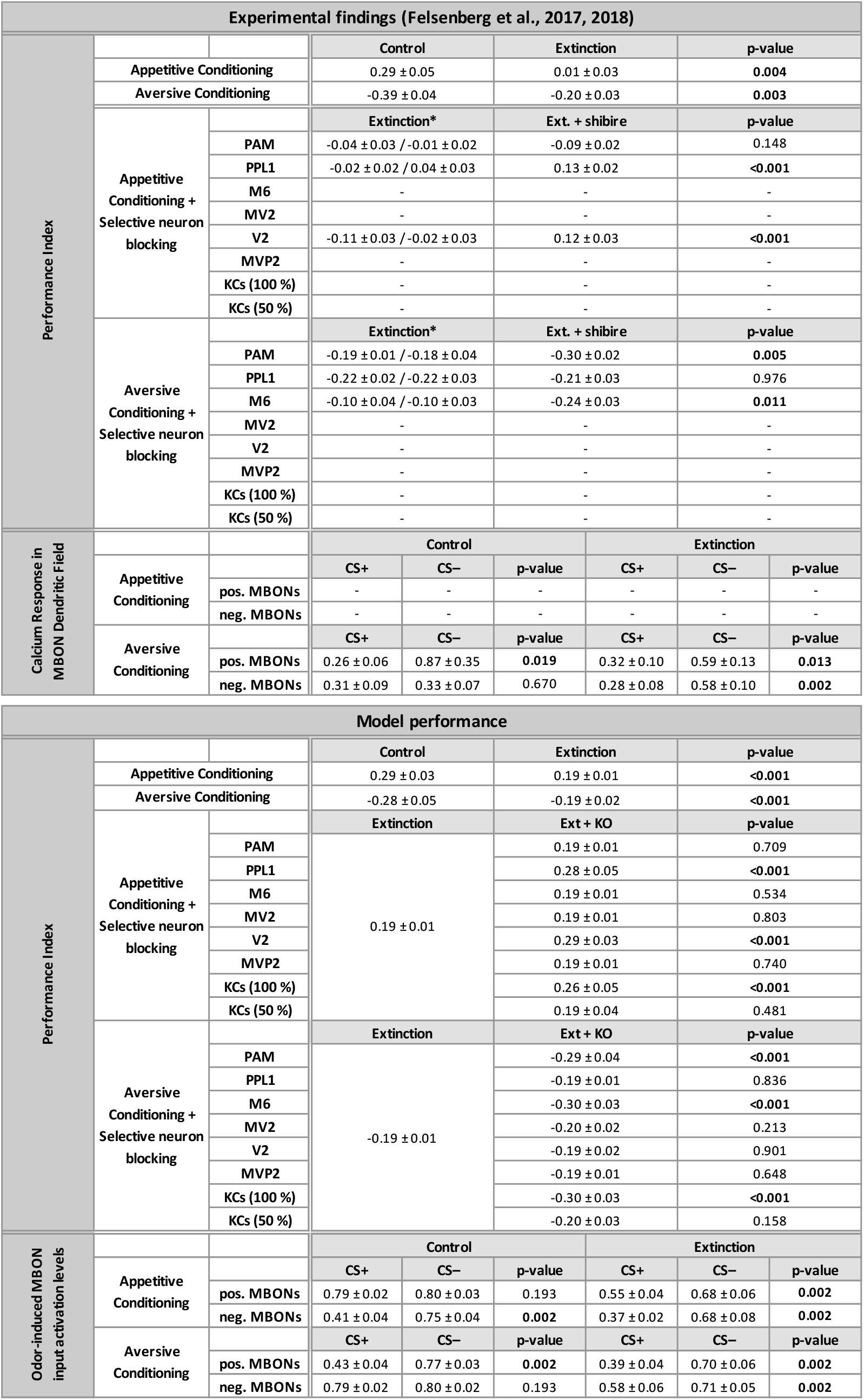
Comparison between experimental and model results. The experimental data on appetitive and aversive conditioning is extracted from Felsenberg et al. (2017) and Felsenberg et al. (2018), respectively. For statistical analysis of the simulated data see Methods. Experimental values are shown as mean ± SEM, model values are shown as mean ±STD. Significant p-values are set in bold font. *The behavioral data include two control groups, the first carried the GAL4 gene only, the second the UAS gene only. The test group carrying both, the GAL4 and the UAS gene, showed cell-specific expression of the temperature-sensitive UAS-*Shibire*^ts1^ transgene.

Repeated reactivation of the CS+ odor alone after either appetitive or aversive conditioning resulted in memory extinction (Figure 3B, C), i.e. in a significant reduction of the learned CS+ approach or avoidance. In addition, generalization to the CS-odor was abolished (Figure 3B, C). The observation that learning is induced when the expected reward or punishment is omitted is in line with the idea of prediction error dependent learning (see Discussion). For a quantitative comparison with the behavioral results of the binary memory test in Felsenberg et al. (2017) we computed the performance index of the model by subtracting the preference index for CS-from the preference index for CS+ (see Methods). The appetitive conditioning protocol in the model and *in vivo* yielded similar performance indices (model: 0.29±0.03 mean ± STD, experiment: 0.29±0.05 mean ±SEM). Memory extinction led to a significantly reduced approach performance (0.19±0.01, Figure 2B). However, the effect of memory extinction in the behavioral experiment was considerably stronger, effectively erasing the initial behavioral memory expression (0.01±0.05, Felsenberg et al., 2017). The aversive conditioning protocol led to a negative performance index of −0.28±0.05 (mean ± STD) for the model, which in absolute terms was smaller than the experimental value of −0.39 ± 0.04 (mean ± SEM, Felsenberg et al. 2018). The reactivation of the CS+ odor after the aversive conditioning led to a significantly reduced avoidance behavior (−0.19±0.02), very similar to the experimental results (−0.20±0.03). A detailed quantitative comparison of experimental and model results is provided in Table 1.

### Memory is rapidly established within a single trial

As a next step, we investigated the dynamics of plasticity during the course of learning and extinction. To this end we monitored the activation rates of DANs and MBONs and the gross synaptic input to the MBONs across trials during single network simulations for appetitive (Figure 4A, B) and aversive (Figure 4C, D) conditioning and extinction.

**Figure 4:**
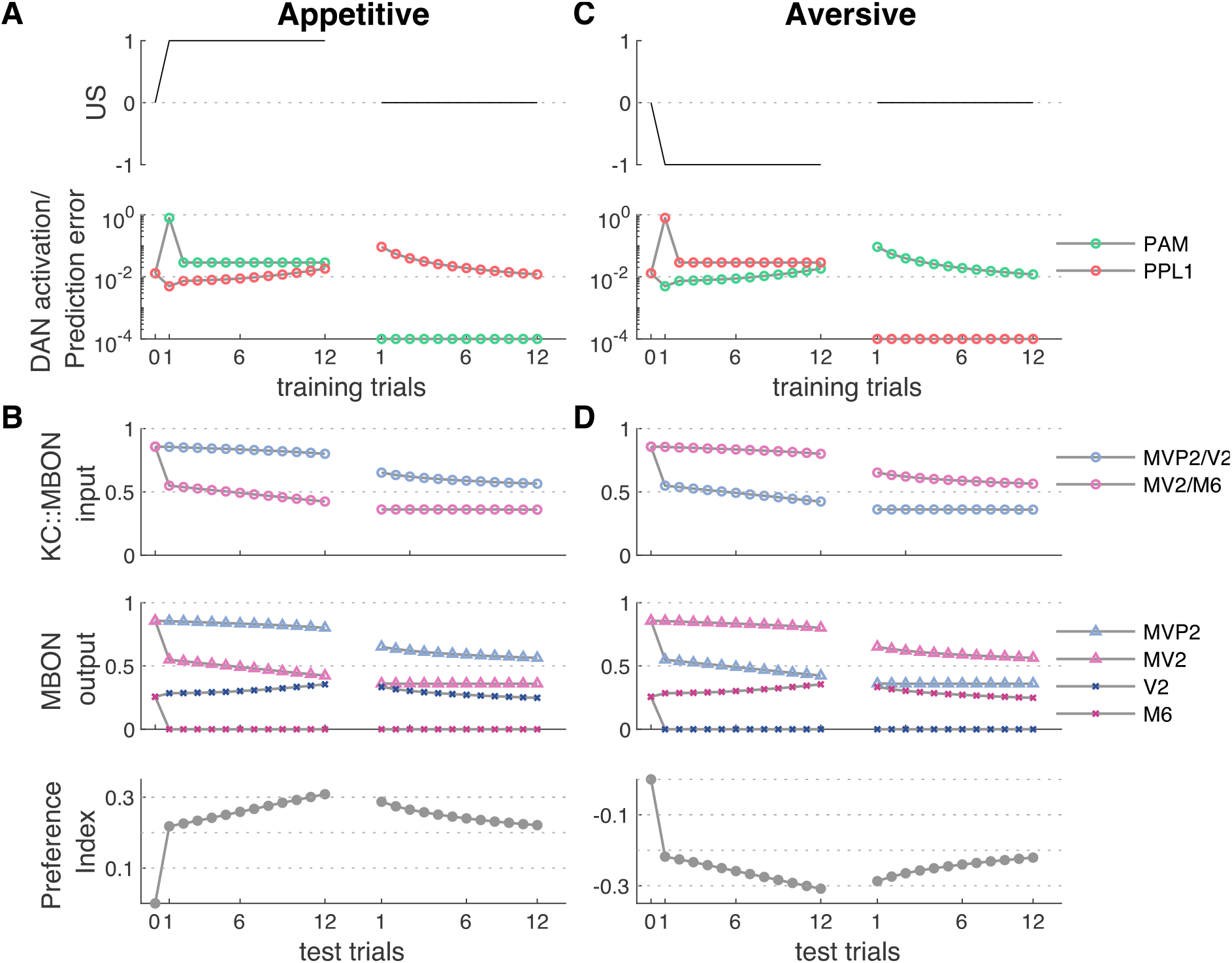
Neuron activation rates and behavioral performance across trials. **A)** Reward value (US, top) and DAN activation rates (bottom) across 12 initial conditioning trials (left) and 12 extinction trials (right). Trial zero indicates neuron activation rates in the naïve model before learning. Reward presentation resulted in a prominent peak activation of the PAM during the very first training trial, i.e. when the actual reward strongly deviates from the predicted reward. Omission of the reward during the first extinction trial (right) resulted in a step-like reduction of PAM activation the simultaneous increase in PPL1 activation, which is crucial for extinction learning. **B)** Summed synaptic KC input to MBONs (top), MBON activation rates (middle), and model preference index (bottom) for each simulated test trial that followed the respective training trial (see Methods). The strong PAM activation during the first conditioning trial resulted in a strong change of the synaptic weights between KCs and MV2 and M6, and consequently in a step-like reduction of the synaptic input and the output activation rates. Consequently, the model preference index represents the imbalance between M6 and MV2 activation rates showed a step-like increase predicting a switch-like expression of a conditioned response behavior after only a single conditioning trial. Additional training led to a further gradual reduction of gross synaptic input to and activation of MV2 and M6, paralleled by the gradual increase of the preference index. Extinction learning led to a gradual reduction of synaptic weights in the KC:MVP2 and KC:V2 pathway. This reduces the difference between M6 and MV2 activation and leads to a gradual extinction of the preference index. **C,D)** Same as A and B for aversive conditioning (left) and subsequent extinction (right). Simulation results are shown for a single network. This was initiated identically before appetitive (A,B) and aversive (C,D) conditioning to enforce identical initial conditions stressing the symmetric mechanism of reward prediction.

In both, appetitive and aversive conditioning, the first pairing of CS+ and US induced a strong response in either the PAM or PPL1, respectively. This strong neuromodulatory signal resulted in a significant reduction of the respective KC::MBON synapses and led to a strongly reduced KC::MBON input to either MV2/6 or MVP2/V2 during the second training trial. As a consequence, feedback excitation from M6 to PAM (V2 to PPL1) in appetitive (aversive) learning is abolished from the second trial onward, leading to further synaptic weight modulations that are weak compared to those in the very first training trial. The evolution of plasticity is reflected in the model performance index, indicating a switch-like increase after the first training trial and moderate but steady increase after subsequent trials. Saturation of the learning effect becomes visible only after ~10-12 trials (cf. Figure 2B). In conclusion, the model learns within a single trial and more training leads to a stronger valence signal as encoded in the imbalance of MV2 and MVP2 activation.

Extinction learning follows a different and slower dynamics in our network model. While already a single presentation of the CS+ odor without reinforcer leads to the induction of the extinction memory and a significant reduction of the performance index (Figure 4), we observe an only gradual extinction that saturates across the 12 extinction trials used here. No complete extinction is observed in our model.

### Associative learning and extinction learning establish two separate memory traces

We tested the hypothesis that associative learning and subsequent extinction learning form two parallel and distinct memories at the physiological level. To this end we mimicked experiments in Felsenberg et al., (2018), in which the authors measured KC dendritic input to individual MBONs by means of *in vivo* calcium-imaging from the dendritic field.

We first consider the case of aversive conditioning and subsequent extinction learning. After initial conditioning, the CS+ induced synaptic input to the MVP2 and V2 neurons is significantly lower than the CS-induced input (Figure 5C). The synaptic input to the M6 and MV2 neurons, however, is similar for both odors (Figure 5D). The extinction protocol, i.e. the subsequent reactivation with CS+ alone (Figure 3A), did not alter KC synaptic input to MVP2 and V2 (Figure 5C). In contrast, CS+ induced synaptic input to M6 and MV2 was significantly reduced after extinction and in comparison to the CS-induced input (Figure 5D). These results are fully consistent with the *in vivo* calcium-imaging results by Felsenberg and colleagues (Felsenberg et al., 2018); cf. Table 1) who observed a significantly reduced dendritic calcium activation in MVP2 in response to the CS+ stimulus after aversive conditioning, while there was no significant change in M6 activation (Figure 3 in Felsenberg et al., 2018). After memory extinction, the decreased CS+ response in MVP2 remained while additional plasticity was observed as CS+ specific reduction in calcium levels in M6.

**Figure 5:**
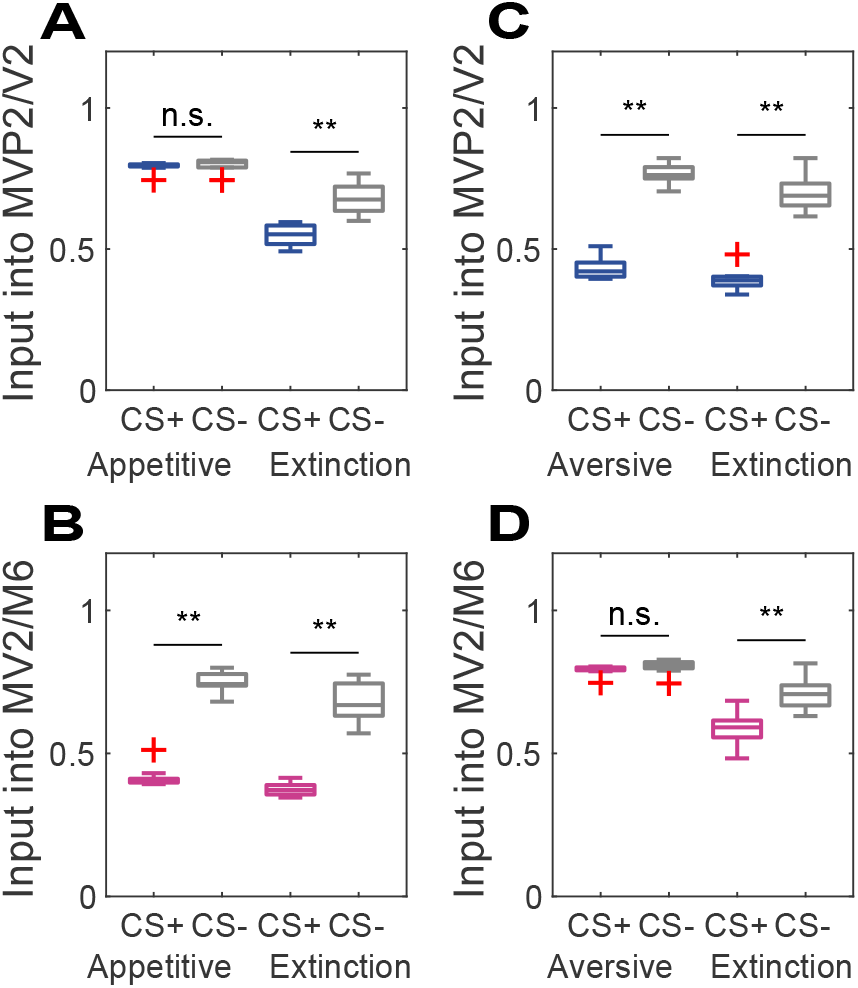
Two separate memories underlie initial and extinction learning. CS+ odor induced activation rates of MBONs were measured before and after extinction training in order to locate associative and extinction memories. **A, B)** Appetitive learning led to a relative depression of CS+ odor induced activation rates of MV2 and M6, but not MVP2 and V2 MBONs. The memory trace in MV2/M6 stayed intact even after appetitive memory was extinguished. Extinction decreased the CS+ response of MVP2/V2, establishing a parallel memory trace. **C, D)** Aversive conditioning led to a reduced CS+ input into MVP2/V2, but not into MV2/M6. After memory extinction, the memory trace in MVP2/V2 remained and we observed an additional decrease in CS+ response in MV2/M6. Results across simulation of n=10 networks in all panels.

Our circuit model predicts a symmetric network behavior for the extinction of an appetitive memory. After appetitive conditioning, the CS+ induced excitatory input was lower than the CS-mediated input to the avoidance-mediating M6 and MV2 (Figure 5B). However, the input to the approach mediating MVP2 and MV2 remained unchanged (Figure 5A). Reactivation of the memory by stimulation with the CS+ odor alone led to a reduction of the excitatory dendritic input into MVP2 and V2 (Figure 5A). As a result, the CS+ preference and performance indices acquired during initial conditioning were significantly reduced after memory extinction (Figure 3B, D).

### Distinct pathways are crucial for appetitive and aversive memory extinction

To validate the idea of recurrent feedback as a teaching signal in our model we tested which neurons are necessary for extinction learning. To this end we mimicked shibirets1 experiments performed by Felsenberg et al., (2017, 2018) who selectively suppressed the activity of individual neurons or neuron clusters during CS+ odor reactivation (see Methods, Figure 6).

**Figure 6:**
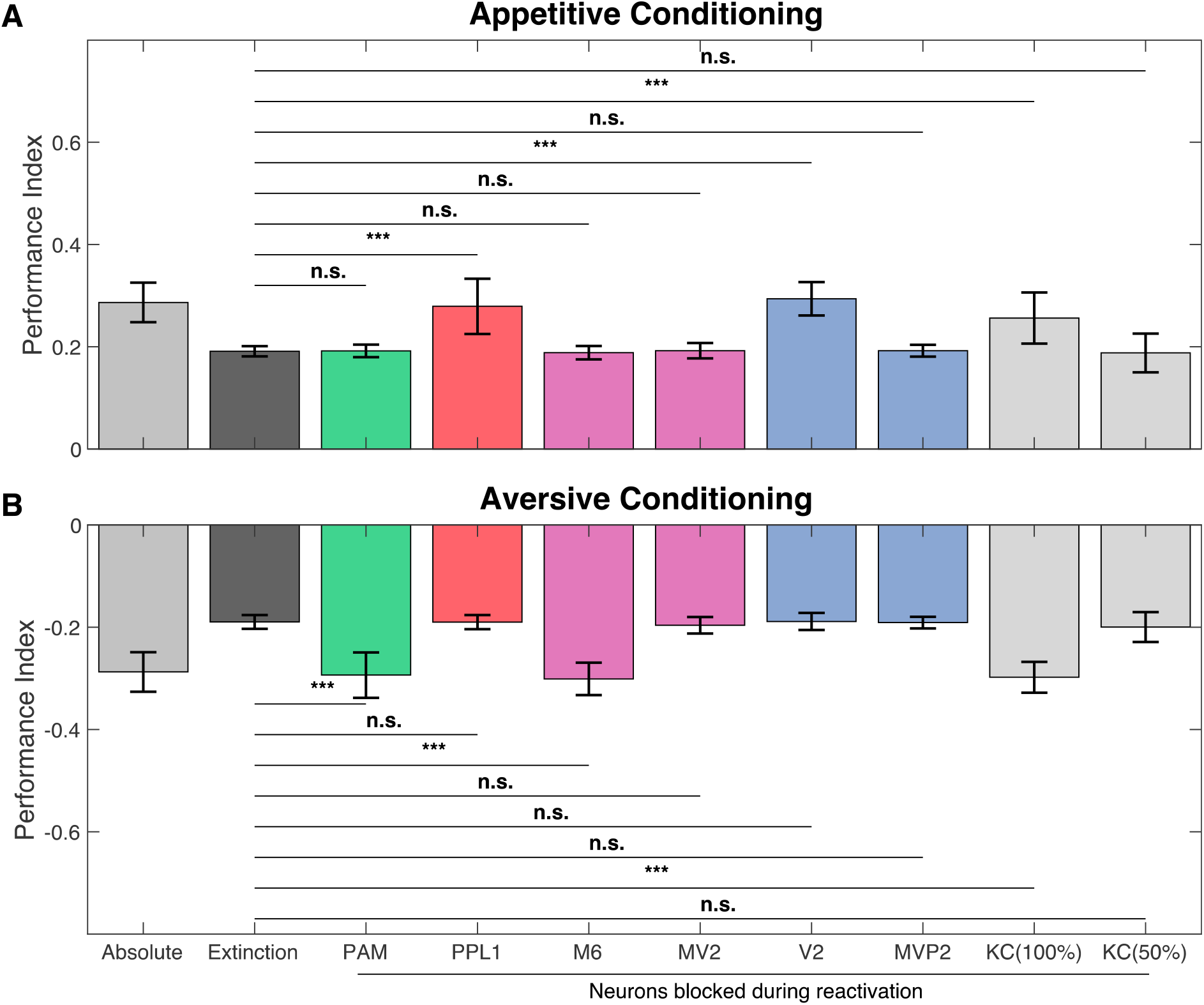
Blocking specific neuron groups can reverse memory extinction. The computational model was utilized to reproduce the neurogenetic manipulations with temperature-sensitive shibirets1 mutant flies. The activation of individual neurons was suppressed during extinction training only. This allows to identify neurons that are crucial to the process of establishing the extinction memory in our model. **A)** Knocking down PPL1 or V2 during extinction training led to a completely abolished the appetitive extinction memory. Blocking PAM, M6, MV2 or MVP2 neurons during odor re-exposure had no effect on the appetitive extinction memory. Blockage of all KCs (100%) prevented the model from extinction learning. However, when only 50% of the KC population was blocked during odor reactivation extinction learning remained unaffected. **B)** Suppressing the activity of PAM DANs, M6 neurons or KCs (100%) during CS+ odor re-exposure after aversive conditioning prevents the model from forming the aversive extinction memory. However, blocking PPL1 DANs, MV2, V2, MVP2 or KCs (50%) does not diminish the memory extinction. Data is presented as mean ± STD; n=15.

We first tested extinction of appetitive memory. Selective blocking of PPL1- or V2-cluster during the reactivation phase abolished the extinction of an appetitive memory. In contrast, blocking of PAM had no effect on extinction (Figure 6A). These findings agree with the experimental results (Table 1) in Felsenberg et al., (2017). Conversely, in the extinction of aversive memory, Felsenberg et al., (2018) showed that the behavioral effect of extinguishing the appetitive memory was significantly diminished by either blocking the PAM cluster or the M6 neurons. However, blocking the vesicle release machinery of the PPL1 cluster had no effect on aversive memory extinction (Table 1). Our model showed the same qualitative result where blocking of either the M6 or the PAM but not of the PPL1 neuron abolished the extinction of an aversive memory (Figure 6B).

To formulate novel predictions for future experiments we further analyzed our model by silencing other MBONs during the reactivation phase (Figure 6, Table 1). Deactivating V2 during the reactivation phase did not significantly alter the aversive extinction memory. Conversely, a blockage of M6 during the extinction process had no effect on the appetitive extinction memory performance either. Blocking MVP2 or MV2 during odor reactivation had no effect on both, appetitive and aversive memory extinction. Thus, our model predicts that a single MBON, V2 or M6, is exclusively involved in the formation of the appetitive or aversive extinction memory, respectively.

### KC activity is partly required for extinction learning

We additionally investigated the role of KCs in extinction learning in our model. To this end, we applied the shibirets1 protocol to KCs. When we blocked all KCs during the training phase, neither appetitive nor extinction memory could be established. However, when we randomly chose and blocked 50% of all KCs during appetitive and aversive extinction training, we did not observe an effect on extinction memory acquisition (Figure 6). Thus, partially blocking KC activity during odor reactivation might not or only mildly interfere with extinction learning while an almost complete block of all KCs should prevent extinction learning in the fly (Supplemental Figure 1, see also Discussion).

## Discussion

In the present study we established a circuit model of the fruit fly based on confirmed anatomy and physiology. It can explain formation and extinction of an olfactory memory, and single trial learning. We discuss the implications, limitations and predictions of our model.

### Prediction error coding

The prediction error theory hypothesizes that learning takes place if there is an unexpected change in the valence of a stimulus (Rescorla & Wagner, 1972). The model performance index in Figure 4 computed after each training trial mimics behavioral performance across trials and fits the prediction error theory and experimental studies. The behavioral learning effect is reduced across trials and the learning curve saturates with increased training. The results match the well-known saturation in the conditioned response (CR) behavior in the honeybee (Pamir et al., 2011). This effect, typically observed across a group of animals, has been formalized in the Rescorla-Wagner model (Rescorla & Wagner, 1972). A recent rate-based model of the fly MB (Bennett, Philippides, & Nowotny, 2019) assumed that connections from MBONs to DANs are crucial for asymptotic learning based on reward prediction. Our model proposes that prediction error coding in *Drosophila melanogaster* is achieved through a network mechanism supported by the specific circuit motif sketched in Figure 1. It is suggested that DANs receive relevant information not only from sensory neurons directly but also via odor activated MBONs (Felsenberg et al., 2017, 2018). It was implemented in the model through excitatory connections from laterally connected MBONs to DANs. Initial conditioning of the model leads to activation of the PAM (PPL1) DAN and therefore induces down-regulation of the aversive (appetitive) MBONs. This learning effect declines after the first learning trial, since MBON activity is not sufficient to activate DANs after being downregulated.

### Single trial learning

Learning within a single trial is a fundamental ability of vertebrates (Cook & Fagot, 2009; Irwin, Banuazizi, Kalsner, & Curtis, 1968) and invertebrates, and the underlying mechanisms are at least partially conserved across phyla. In insects, single trial learning has been intensely studied in classical and operant olfactory, tactile, and visual conditioning of the honeybee *Apis mellifera* (Menzel, 1999; Menzel, Erber, & Masuhr, 1974; Pamir, Szyszka, Scheiner, & Nawrot, 2014; Sandoz, Roger, & Pham-Delègue, 1995; B. H. Smith, 1991; Villar, Marchal, Viola, & Giurfa, 2020). Appetitive olfactory conditioning of the proboscis extension response allows the observation of the all-or-none CR behavior during successive training trials where the onset of US presentation is delayed with respect to the onset of the CS odor. Typically 40-60% of bees show a CR after a single pairing of odor with reward (Pamir et al., 2014) and a single learning trial is sufficient to establish short- and long-term memory, which can be recalled up to 3 days after training (Menzel, 1999; Pamir et al., 2014; B. H. Smith, 1991; Villar et al., 2020). Additional training trials lead to a saturation after ~3-4 trials with respect to the fraction of animals that express a CR.

In the fruit fly, single trial learning has been established more recently. Aversive conditioning of adult flies with electric shock punishment showed that pairing of a single odor presentation (of 10-60s duration) with a single electric shock induces short term memory (Beck, Schroeder, & Davis, 2000; Scheunemann et al., 2013) and long term memory, retrievable for up to 14 days if the experimental context is kept strictly constant for training and memory test (Zhao et al., 2019). In appetitive conditioning, a single session of odor-reward pairing could establish long term memory (Colomb, Kaiser, Chabaud, & Preat, 2009; Krashes & Waddell, 2008). Weiglein et al. (2019) could show that also in the fruit fly larva a single session of appetitive conditioning of 5 min was sufficient to establish a short-term memory where the performance index in the memory test increased with increasing duration (in the order of minutes) of the training session.

In our circuit model, the very first training trial induces a switch-like change in the network response and in the behavioral memory expression. This parallels the switch-like behavioral dynamics observed in learner bees (Pamir et al., 2011) and provides an explanation for the single-trial induction of appetitive and aversive memories in the fruit fly (Colomb et al., 2009; Krashes & Waddell, 2008; Zhao et al., 2019). In contrast, during extinction learning we propose that an extinction memory builds up gradually in the opposing pathway.

### Extinction Learning

Experimental studies in the fruit fly have shown that extinction learning establishes distinct and opposing memory traces. The extinction of reward memory requires punishment coding dopamine neurons whereas extinction of aversive memory is mediated by reward coding neurons (Felsenberg et al., 2017, 2018). Recent findings in mice (Salinas-Hernández et al., 2018) and rats (Luo et al., 2018) suggest that this principle for extinction learning is conserved in mammals. The authors showed that dopaminergic neurons from the ventral tegmental area associated with reward signaling are required to extinguish fear memory.

The importance of MBON::DAN feedback in order to perform complex learning tasks such as extinction has been formalized recently in a rate-based model of the *Drosophila* larva (Eschbach et al., 2020). In our adult fly model, the MBON::MBON and MBON::DAN connections for the extinction of aversive and appetitive memories are strictly symmetrical. During extinction of an aversive memory the dopaminergic reward signal of the PAM neuron in extinction trials is driven by excitation from the M6 MBON, which receives reduced inhibition from the approach mediating MVP2 as a result of the initial aversive conditioning that reduced KC drive of MVP2. Thus, extinction learning establishes a reward-like extinction memory trace in parallel to the initial aversive memory trace (Figure 5D). Intriguingly, the initial memory trace is not altered by extinction (Figure 5C). This matches the experimental results and the proposed mechanism in Felsenberg et al. (2018) supporting the long-standing hypothesis of two parallel memory traces after extinction (Bouton, 2004; Dudai, 2004; Eisenhardt & Menzel, 2007).

The strict symmetry of the recurrent pathways in our model has the consequence of symmetrical quantitative results for the performance index and neuron input activation rates (Table 1). The experimental situation, however, does not provide fully symmetric behavioral results. Our model matches well in the aversive memory pathway, i.e. for appetitive conditioning and extinction of aversive memory. In aversive conditioning experiments, however, Felsenberg et al. (2017, 2018) obtained lower performance indices than our model and extinction of the appetitive memory was complete (Table 1). Independent tuning of model parameters for the parallel pathways would allow for an improved quantitative match with all experimental results.

### Limitations of the model

Our network model (Figure 1) can be viewed as a minimal circuit model of the MB sufficient to explain prediction error coding, single trial conditioning and extinction learning. We focused on only four out of 34 MBONs and restricted MB connectivity to feedforward from KCs to four MBONs and feedback from MBONs to DANs but did not explore other synaptic contacts among KCs and between KCs, MBONs and DANs within the MB lobes as uncovered in the fly EM connectome (Takemura et al., 2017). Learning in our model is achieved through reinforcement-mediated plasticity at a single synaptic site, which has been shown to be involved in appetitive and aversive short-term olfactory memory (Owald & Waddell, 2015; Pascual & Préat, 2001; Zars, Fischer, Schulz, & Heisenberg, 2000). Increasing experimental evidence indicates that olfactory learning can involve plasticity at multiple sites within the insect MB (Bouzaiane et al., 2015; Haenicke, Yamagata, Zwaka, Nawrot, & Menzel, 2018; Kremer et al., 2010; Pascual & Préat, 2001; Trannoy, Redt-Clouet, Dura, & Preat, 2011; Yamazaki et al., 2018). Also, it has been shown that memories can co-exist on different time scales representing short-, mid-, and long-term memories (Davis, 2011; Pascual & Préat, 2001; Trannoy et al., 2011; Yamagata et al., 2015). Our trial-based model approach does not accommodate explicit time scales that would allow to differentiate between memory acquisition and consolidation (Felsenberg et al., 2017) or decay (Shuai et al., 2015). Future extensions may retain full temporal dynamics, e.g. by using spiking neural network models that have previously been used successfully to study classical conditioning in fruit flies (e.g. Faghihi, Moustafa, Heinrich, & Wörgötter, 2017; Gupta, Faghihi, & Moustafa, 2018; Rapp & Nawrot, 2020; D. Smith, Wessnitzer, & Webb, 2008; Wessnitzer, Young, Armstrong, & Webb, 2012).

In the present study we aimed at reproducing the behavioral performance index that is measured in a group forced choice paradigm (Quinn, Harris, & Benzer, 1974) and thus represents a population measure across individuals, each performing either the correct or the incorrect behavioral choice. The long-held notion states that expression of the CR behavior in individual flies is stochastic and follows the group-averaged behavior (Quinn et al., 1974) has been questioned by Chabaud, Preat, & Kaiser (2010) and by studies in the honeybee (Pamir et al., 2011,2014; Haenicke et al., 2018) and in the cockroach (Arican et al., 2020). They have shown that the average CR in a group of animals does not accurately reflect memory expression in individuals. Rather, it confounds two subgroups of learners and non-learners where learners showed a switch-like and stable expression of the CR behavior mostly after a single trial (average 1,7 trials) with high memory retention rates (>90%). Different parameters may account for individual learning abilities such as internal state e.g. of hunger and satiety (Pamir et al., 2014; Sayin, Boehm, Kobler, de Backer, & Grunwald Kadow, 2018) that could influence perception or motivation. Establishing neural circuit models that take into account individuality of learning and individual parameters that influence memory formation is a challenge we aim to address in future model studies.

### Model predictions

Our model makes several predictions that can be tested experimentally. First, we hypothesize that DANs in the PAM and PPL1 cluster compute a prediction error. Specifically, in naïve animals and for the example of appetitive learning the PAM should show spontaneous baseline spiking activity and no stimulus response during the presentation of a novel odor. During a first pairing of the CS+ odor with a reward (US) the PAM cluster should show a clear odor response, which will be strongly reduced after the first learning trials (Figure 4A). When reward is omitted during extinction trials or in the memory test, we predict a response in which the PAM reduces its spiking output below spontaneous level. This prediction matches the observation in dopaminergic neurons in the monkey (Schultz et al., 1997) and in the octopaminergic VUMmx1 neuron in the honeybee (Hammer, 1997; Montague, Dayan, Person, & Sejnowski, 1995), although the circuit mechanisms hypothesized here differs from that proposed by Schultz et al. (1997) and Hammer (1997).

Second, we hypothesize a switch-like induction of a memory trace during a single training trial, implying a step-like change in the MBON activation rates (Figure 4). A second or third training trial should only induce gradual changes in cellular physiology. Likewise, the behavioral memory expression should be observable after a single learning trial.

Third, our systematic analysis of blocking experiments (Figure 6) for all MBONs indicate that MV2, M6 and MVP2 do not play a role during appetitive memory extinction, whereas MVP2, MV2 and V2 are dispensable for aversive extinction learning. This could be experimentally tested by blocking these MBONs specifically during extinction trials only. While the co-existence of two memory traces after extinction of an aversive memory has been shown in Ca-imaging from MVP2 and M6 (Felsenberg et al., 2018, Fig. 5D) our model predicts parallel memory traces also for extinction of an appetitive memory (Fig. 5A,B).

Finally, in a study by Schwaerzel et al. (2002) in which KCs were blocked during aversive conditioning and odor reactivation, the authors argued that KC activity is dispensable for the acquisition of an extinction memory. However, the two GAL4 lines used in this study include less than 50% of all KCs (approx. 700 and 850 cells). Our model analysis predicts that a sizable fraction of the KC population is required and sufficient to form an extinction memory (Figure 6, Supplemental Figure 1). A complete block of KCs however would interrupt the PN::KC::MBON::DAN signaling pathway and extinction memory could not be acquired. This could tested in a future blocking experiment using e.g. the OK107-GAL4 line that has been reported to label practically all KCs (Aso et al., 2009).

## Methods

### Circuit architecture

Our circuit model for olfactory coding and olfactory memory formation (Figure 1) consists of three neuron layers (PNs, KCs, MBONs) representing the three major stages of the olfactory pathway in *Drosophila* and two reinforcement mediating dopaminergic neurons representing the PAM cluster. Each neuron in the circuit can assume an activation rate in the range of 0 to 1.

Olfactory model input was simulated through the activation of 50 out of the total 100 PNs. This matches experimental observations of about 40%-60% of PNs being activated by a single odor stimulus (Krofczik et al., 2009; Brill et al., 2013; Wilson, 2013) and follows the model of Peng & Chittka (2017). Each PN is activated with a random rate drawn from a uniform distribution in the range between 0.2 and 0.8. PNs are connected to 2000 KCs via the weight matrix W1 where each PN is connected to 5-15 KCs, each connection has a fixed synaptic weight of 0.2. Activation of the KC vector in the next layer resulted from the matrix product of the PN population vector and the respective weight matrix W1 according to

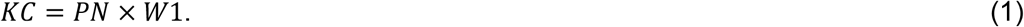

Out of the total 2000 KCs only the subpopulation of 100 KCs (5%) with the highest activation rate kept its activation. All other KCs are set to zero to enforce population sparseness (Peng & Chittka, 2017). In a next step, KCs are fully connected to the four MBONs via the weight matrix W2, with all synaptic weights initially set to 0.01. The excitatory input to each MBON was calculated as the matrix products

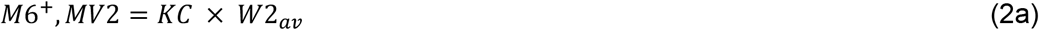

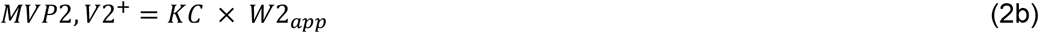

We further included lateral connectivity between the MBONs. The M6 MBON receives inhibitory input from MVP2, whereas V2 gets inhibited by MV2. The respective inhibitory inputs are formalized according to

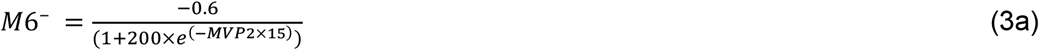

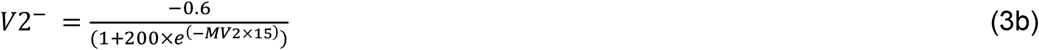

The activation rate of M6 and MV2 results from a summation of inhibitory and excitatory input as

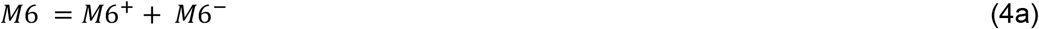

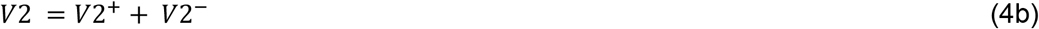

whereas for MVP2 and MV2 the activation rate is solely determined by the excitatory input.

The PAM and PPL1 DAN receive excitatory input by the M6 and V2 neurons, respectively. Additionally, reinforcing stimuli have an effect on both DANs. A rewarding unconditioned stimulus (US = +1, positive reinforcer) leads to an excitatory input to the PAM (Eqn. 5a). At the same time, excitatory input from the V2 to the PPL1 is partially suppressed (represented by the factor ρ = 0.8 in Eqn. 5b). Conversely, a punishing unconditioned stimulus (US = −1, negative reinforcer) results in excitation of the PPL1 (Eqn. 5b) and in the suppression of excitatory input from M6 to the PAM (Eqn. 5a). The total DAN input was thus computed as

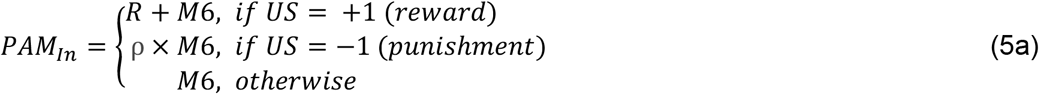

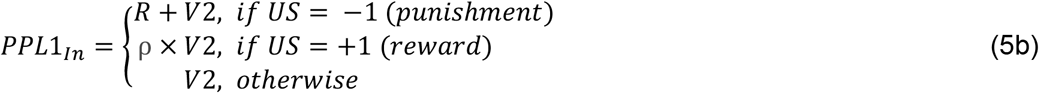

We fixed the US induced excitation to R = 0.3 for all our experiments. However, generally the parameter R allows for a modeling of variable reward magnitudes linked to the US such as e.g. different levels of sugar concentration contained in a rewarding US.

The output activation rate of each DAN encoding the prediction error is calculated with a sigmoid transfer function

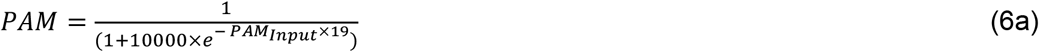

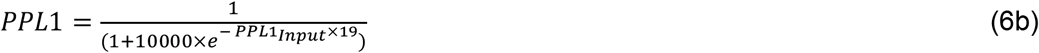

### Plasticity rule at KC::MBON synapses

We implemented synaptic plasticity at the KC::MBON synapses. In the initial state of the model, the weights of all KC::MBON synapses are set to the same fixed value of 0.01. Each of the synaptic weights in W2 is subject to synaptic plasticity. At the end of a given trial *t* and for any synapse KC_i_::MBON_j_ a change of the synaptic weight w_ij_ occurs if both, the presynaptic KC and the respective DAN were active in trial *t*. If both conditions are met, the synaptic weight is reduced according to the two-factor learning rule

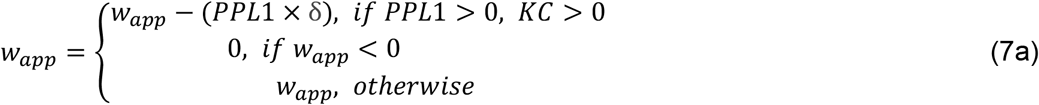

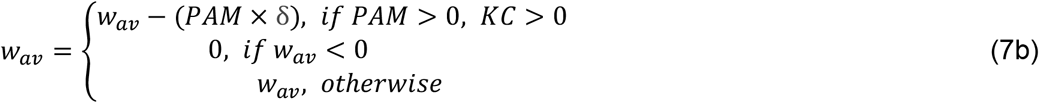

where w_app_ refers to the weight of a synapse onto M6 or MV2 and w_av_ refers to the weight of a synapse onto MVP2 or V2. Note that the weight changes are proportional to the activation rate of the respective DAN. We fixed the learning rate as δ=0.0045 for all our experiments.

### Experimental protocols

We subjected our model to a set of conditioning and extinction protocols. Further, it was operated and evaluated in a trial-resolved fashion where, in each trial, the excitatory and inhibitory synaptic input to each of the neurons and its output activation rate was computed. Within-trial neuronal dynamics were neglected.

#### Classical conditioning

Our classical conditioning protocol consisted of a training phase and a subsequent memory retention test (Figure 2A). During each training trial an odor pattern (CS+) was paired with a positive or negative reinforcement (US). Subsequently, a second odor pattern (CS–) was presented without reinforcement. During retention, we presented the CS+ and CS-stimulus once. The training procedure derives from a group assay developed by Tully & Quinn, (1985) that is often used to study olfactory learning in flies. A group of flies are trained in a Tully-machine with a training arm and two testing arms. In the training arm, the animals are typically exposed to the CS+ in combination with either sugar (reward) or a train of electric shocks (punishment). Subsequently, animals are exposed to the CS-without reinforcer. After the training, the flies are transferred to the testing chamber in which they can choose between a CS+ and a CS– perfused arm. In our standard training procedure, we fixed the number of training trials to n=12. In Felsenberg et al., (2018) the CS+ is presented for 1 minute and combined with 12 electric shocks. The trial-based classical conditioning protocol is a standard procedure in other insect models for learning and memory, notably the honeybee (Bitterman, Menzel, Fietz, & Schafer, 1983; Pamir et al., 2011). For our initial model analysis, we varied the number of training trials in the range of n= 1, …, 24 to quantify the associative strength as a function of n (Figure 2).

#### Extinction learning

The extinction protocol included an additional reactivation phase after training and before memory extinction. During reactivation, the CS+ odor was presented alone, i.e. without reinforcement, during 12 reactivation trials. This procedure again mimics the experimental protocol used by Felsenberg et al. (2018), in which the authors presented the CS+ without shocks for the same duration (1 min) as in the training.

#### Neurogenetic manipulation experiments

To investigate the role of individual neurons in the memory extinction process we selectively suppressed their activation by setting their activation rate to zero during the reactivation phase. Thereby, we mimic neurogenetic manipulation experiments that suppress the activation of specific neurons. We specifically compare our model results to the experimental results performed in Felsenberg *et al.* (2018) based on the neurogenetic tool shibirets1 and we generate experimentally testable hypotheses in novel model experiments.

### Evaluation of model performance and quantitative model predictionsxgo

We evaluated model outcome by different measures that allow for a quantitative comparison with the outcome of animal experiments, both at the behavioral and at the physiological level. Applying these measures in untested experiments allowed us to formulate experimental predictions for novel experiments.

#### Behavioral output

To this end we computed two quantities that can be interpreted as measures of approach or avoidance behavior. We used the activation rates of these MVP2 and MV2 to calculate the *preference index* as

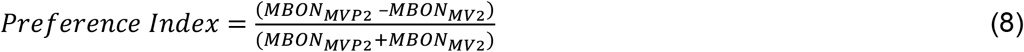

The innate *preference index* of our model is zero by construction.

In behavioral extinction learning experiments (Felsenberg et al., 2017, 2018). The number of animals that chose the CS-odor was then subtracted from the number of animals that chose the CS+ and the result was normalized by the total group size to compute the *performance index* (Tully & Quinn, 1985). In analogy, we calculated a model *performance index* computing the difference between the preference indices for CS+ and CS– as

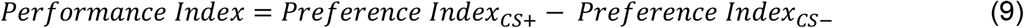

#### Dendritic input

We quantified activation rates of single neurons. Activation rates were measured in both DANs and MBONs. DAN activation rates were obtained during training trials, whereas the MBON activation rates were measured during the test trials (Figure 4). Further, we compared the model to a recent study in which dendritic MBON activity was measured (Figure 5, Felsenberg et al., 2018). KC::MBON synapses in M6 populate the dendritic tree while the MVP2::M6 are located proximal to the dendritic root (Felsenberg et al., 2018). Measuring the calcium activation across the dendritic field can thus be interpreted as the overall excitatory KC input to the MBON. In analogy to the experimental approaches, not only the total synaptic input, but also the sum of excitatory synaptic input from KCs was measured.

#### Performance at the group level

Each new initialization of a network involved the random generation of the odor induced PN activation rates (ranging between 0.8 and 1, evenly distributed) and a random number of PN::KC connections (in the range 5-15, evenly distributed) for each KC to establish variability across individuals.

To test statistical significance of group differences in the performance index, the Wilcoxon rank-sum test was used. To test for differences of MBON activation rates in the same group before and after extinction learning, the Wilcoxon signed-rank test was used. Model results were compared to the experimental results obtained by Felsenberg et al. (2017, 2018). To extract single data points from the experimental studies, the GRABIT tool by MathWorks was applied. Model simulation and statistical analyses were performed with MATLAB_R2018b (Mathworks Inc, Natick, MA). The full code for the MB model is available on the Github account of the Nawrot lab: https://github.com/nawrotlab

## Acknowledgements

We are grateful to Johannes Felsenberg who has been supporting this project throughout with his expertise and fruitful ideas. We thank Anna-Maria Jürgensen for her contribution to the initial model implementation. We thank Bertram Gerber and Anna-Maria Jürgensen for insightful comments on an earlier version of this manuscript.

---

**Supplemental Figure 1:**
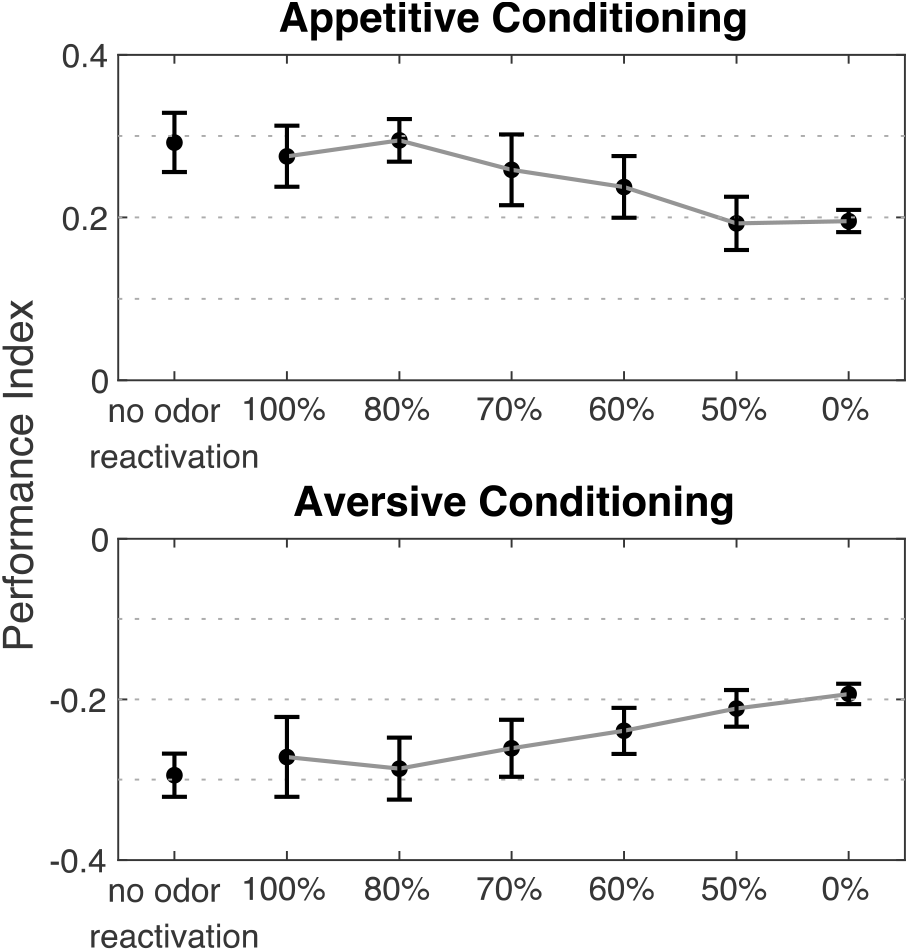
The percentage of blocked KCs during CS+ odor reactivation determines the effect on extinction memory formation. When all KCs are blocked during appetitive or aversive extinction training no extinction memory is formed. However, reducing the percentage of blocked KCs gradually increases the ability to establish extinction memory. When 50% or less KCs are blocked, memory extinction formation is not impaired. Data is presented as the mean ± STD; n=15.

